# Left-handed DNA-PAINT for improved superresolution imaging in the nucleus

**DOI:** 10.1101/2020.03.28.010553

**Authors:** H.J. Geertsema, G. Aimola, V. Fabricius, J.P. Fuerste, B.B. Kaufer, H. Ewers

**Author notes:** Correspondence to: Helge Ewers, Thielallee 63, 14195 Berlin, Germany.

## Abstract

DNA point accumulation in nanoscale topography (DNA-PAINT) advances super-resolution microscopy with superior resolution and multiplexing capabilities. However, cellular DNA may interfere with this single-molecule localization technique based on DNA-DNA hybridization. Here, we introduce left-handed DNA (L-DNA) oligomers that do not hybridize to naturally present R-DNA and demonstrate that L-DNA PAINT has the same specificity and multiplexing capability as R-DNA PAINT, but greatly improves specific visualization of nuclear target molecules.

---

Novel super-resolution microscopy techniques facilitate the resolution of cellular structures down to the molecular detail. In single molecule localization superresolution microscopy (SMLM) techniques, nanometer resolution is achieved by sequential imaging of a large number of single molecules at nanometer scale resolution, a few molecules at a time^1–3^. Different approaches have been developed to keep the majority of molecules dark while a small fraction can be imaged as single molecules. The commonly used fluorescent proteins and organic dyes are physically or chemically switched between dark and bright states to separate detection events. Recently, DNA point accumulation in nanotopography (DNA-PAINT^4^) was developed as a novel approach that relies on transient hybridization of fluorophore-coupled DNA-oligomer imagers to target-associated reverse-complement DNA-oligomer binders. Since single molecule detection occurs here through DNA hybridization and is uncoupled from dye photophysics, DNA-PAINT allows the use of bright, photostable organic dyes to obtain highest single molecule localization resolution^4,5^. Furthermore, different oligomer sequences enable multiplexing in a single wavelength^4,6^, thereby avoiding chromatic aberration^7^. Finally, the well-understood chemical kinetics of DNA hybridization facilitates quantitative imaging^8,9^. In the light of these advantages, DNA-PAINT is a significant advancement in single molecule localization microscopy.

However, any DNA-nonamer^4^ has statistically on average 22’000 complementary binding sites in the diploid ~ 3 gigabase human genome that can contribute to false positive hybridization events. In addition, the transcriptome may as well contribute to undesired DNA-RNA hybridizations. This is a significant problem in a super-resolution technique based on single molecule localizations, which is one of the most promising approaches to advance our understanding of the nanoscopic functional organization of the nucleus, an area of intensive research in cell biology^10–12^. To overcome this problem, we here employed oligomers synthesized from L-DNA for DNA-PAINT. L-DNA has identical physico-chemical properties to the R-DNA common to life (see Figure 1a and Supp. Fig. 1), but does not naturally occur and cannot hybridize with R-DNA (Figure 1b)^13^. We hypothesized that L-DNA PAINT would perform better in superresolution imaging of nuclear targets than traditional R-DNA-PAINT.

**Figure 1:**
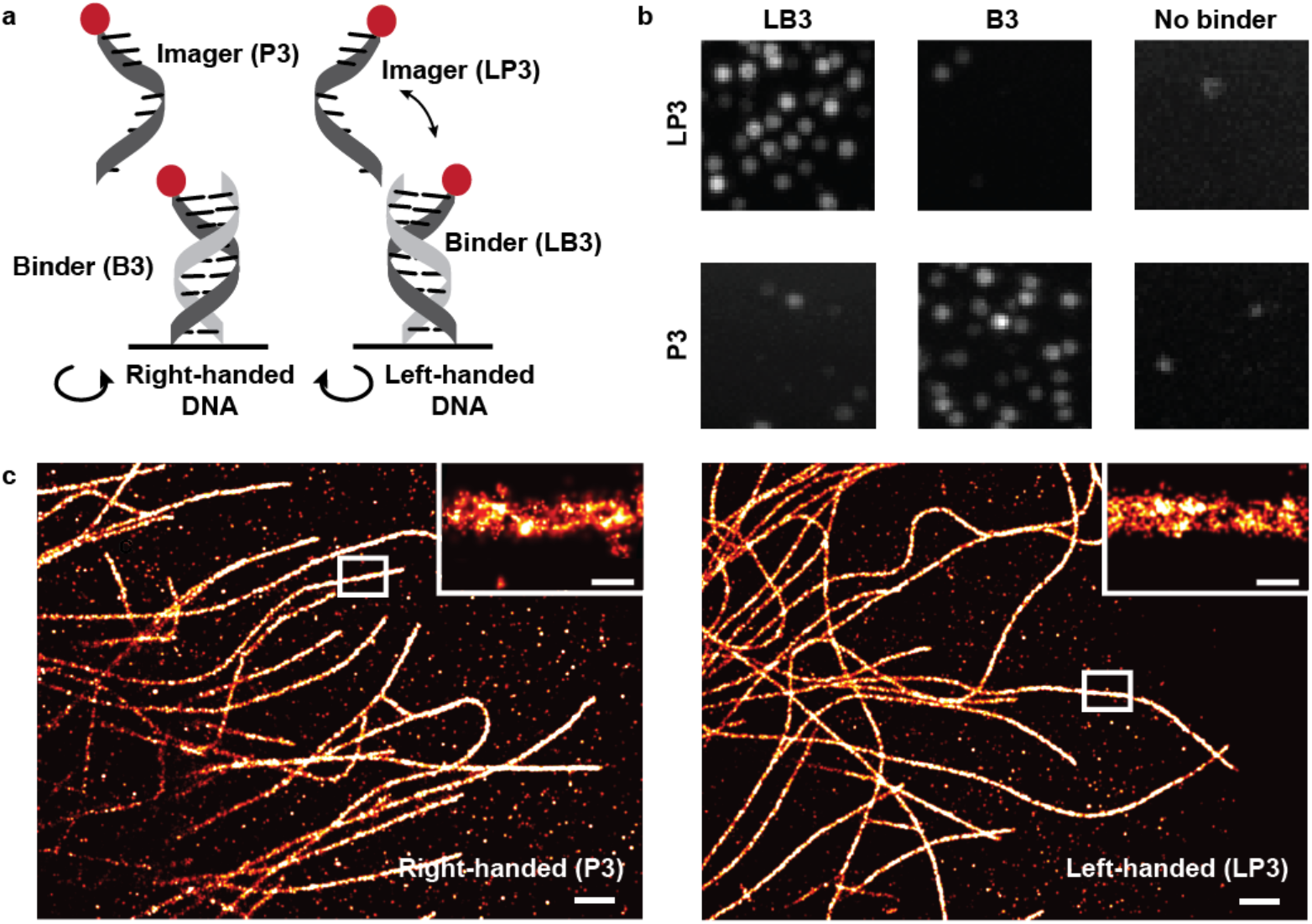
Comparison of DNA-PAINT with left-handed and right-handed DNA oligomers. **a)** Schematic overview of right- and left-handed DNA-PAINT. Transient hybridization events of fluorescence-labeled right- and left-handed DNA imager strands with their respective binder oligomer temporally immobilize them for single molecule localization. **b)** Left-handed (LP3) and right-handed (P3) imager strands were added to surface-immobilized left-handed (LB3), right-handed (B3) binder strands or no binders, respectively. Right-handed imagers were detected only in presence of right-handed binders, left-handed imagers only in presence of left-handed binders. Single molecule images are 5 μm × 5 μm. **c)** Reconstructed DNA-PAINT images generated from right-handed DNA-PAINT (R-DNA-PAINT) and left-handed DNA-PAINT (L-DNA-PAINT) experiments of immuno-stained microtubules in HeLa cells. Insets show representative microtubule segments. Scale bar is 1 μm, scale bar of insets is 100 nm.

We first addressed if the performance of L-DNA-PAINT is comparable to R-DNA-PAINT. To do so, we visualized microtubule structures in HeLa cells and found that R-DNA-PAINT and L-DNA-PAINT performed equally well in resolving microtubules as hollow structures in the cytoplasm, with an average spacing of 39 ± 4 nm (mean ± standard deviation, n = 9) and 33 ± 6 nm (mean ± standard deviation, n = 22) between the two sides of the hollow structure, respectively (Figure 1c), in agreement with previous reports^14^. In addition, the nearest neighboring distance of R-DNA- and L-DNA-PAINT was comparable with 17 nm and 15 nm, respectively. As such, we concluded that L-DNA matches the performance of R-DNA in DNA-PAINT.

Next, we investigated the hybridization potential of R-DNA imagers to genomic DNA by adding fluorescently labeled R-DNA imagers (P3) to fixed Hela cells, in the absence of R-DNA binders. It immediately became apparent, that the localization density (localizations/μm^2^) of fluorescent imagers in the nucleus was higher than in the cytoplasm of the cells (Figure 2a). In contrast, when we subsequently exchanged R-DNA imagers for L-DNA imagers with the same nucleotide sequence (LP3) and in the same cell, localizations in the nucleus were reduced to the background level in the cytoplasm (Figure 2a). Since the localization density of single imagers is highly dependent on the local imager concentration, we represent the ratio of localization density in the nucleus (caused by hybridizations between cellular DNA and imagers) over the cytoplasm (background localizations). We found that detections of the R-DNA imager P3 were significantly enriched in the nucleus (2.5 ± 0.6 (mean ± SEM)-fold, see Figure 2b, c). On the other hand, detections of the L-DNA imager LP3 were not enriched (Figure 2C).

**Figure 2:**
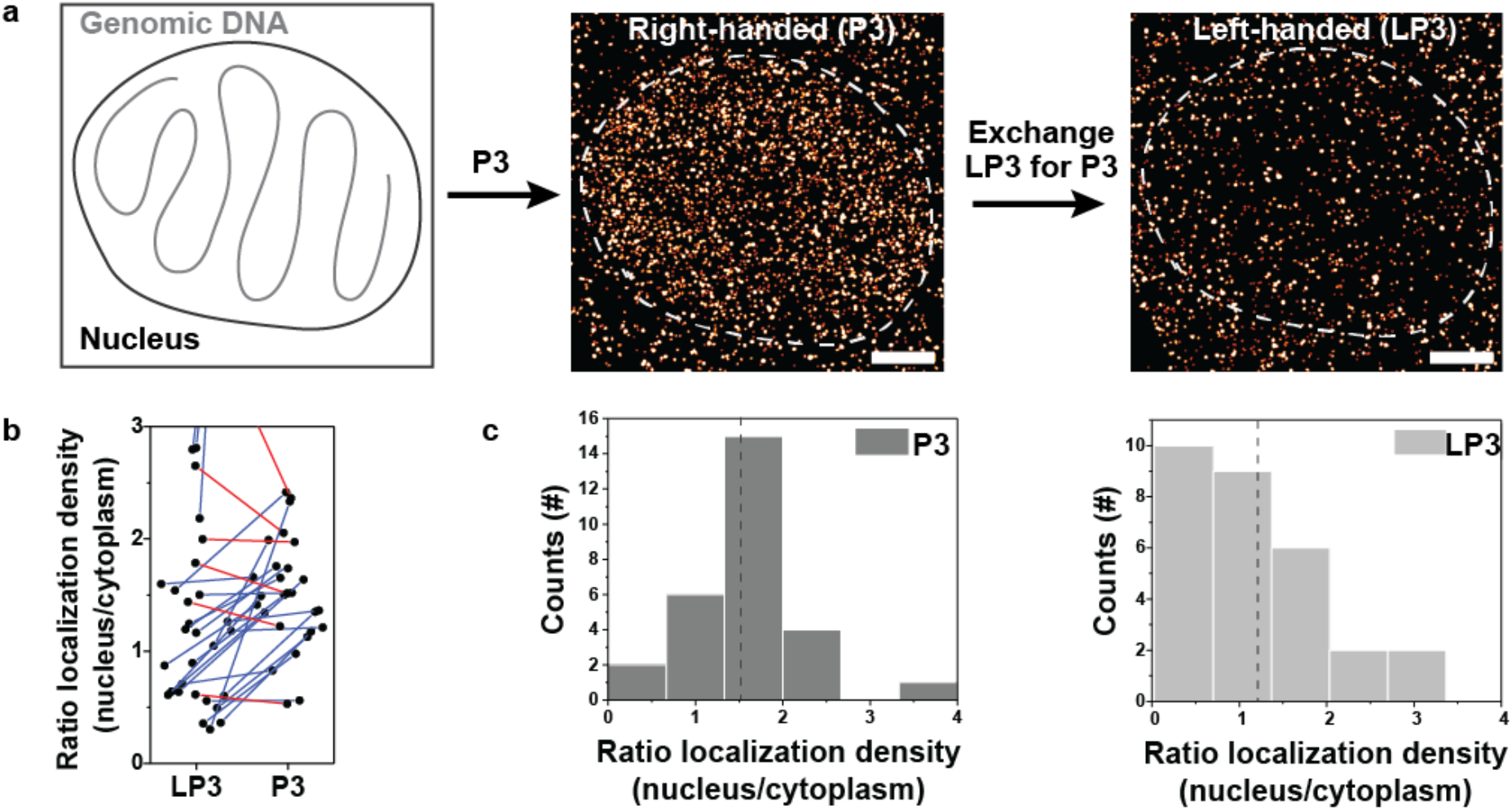
Comparison of nuclear binding of R-DNA imagers and L-DNA imager strands. **a)** Experimental scheme for the detection of non-specific binding of P3 and LP3 imager strands in nuclei of HeLa cells. Fixed cells are imaged in the presence of P3 and then washed and imaged in the presence of LP3 in identical buffer and concentration. Both images are reconstructed from localizations in 5000 frames. Scale bars are 5 μm. **b)** Plot of relative localization densities in the nucleus over the cytoplasm for LP3 vs. P3 (n=30). Lines connect measurements in the same cells for R-DNA and L-DNA, color-coded for higher R-DNA background (blue) and higher L-DNA background (red). **c)** Histograms of the relative enrichment of localizations in the nucleus over the cytoplasm for R-DNA (left) and L-DNA (right). Dashed lines represent median values of 1.5 (R-DNA) and 1.1 (L-DNA).

We expect false positive localizations of R-DNA to be especially detrimental in assays where the cellular DNA is denatured and single DNA strands are quantitatively exposed for hybridization with R-DNA imagers. Such assays are widely used for the investigation of DNA organization in the nucleus. One prominent example is the investigation of DNA replication-sites by the incorporation of the non-natural nucleotide BrdU into the cell’s chromosomes. We performed such an assay and stained incorporated BrdU with antibodies that were coupled to an equal mixture of R-DNA binder (B3) and L-DNA binder (LB3). Cells were imaged first in the presence of 1 nM of P3, washed, and subsequently imaged again in the presence of 1 nM LP3. We found that cells exhibited a prohibitive level of background when incubated with P3, but allowed for specific and super-resolved detection of replication centers in the same cell when incubated with LP3 (Figure 3a and b). We concluded that L-DNA-PAINT enables previously inaccessible experimental assays targeting the cellular replication machinery in cells.

**Figure 3:**
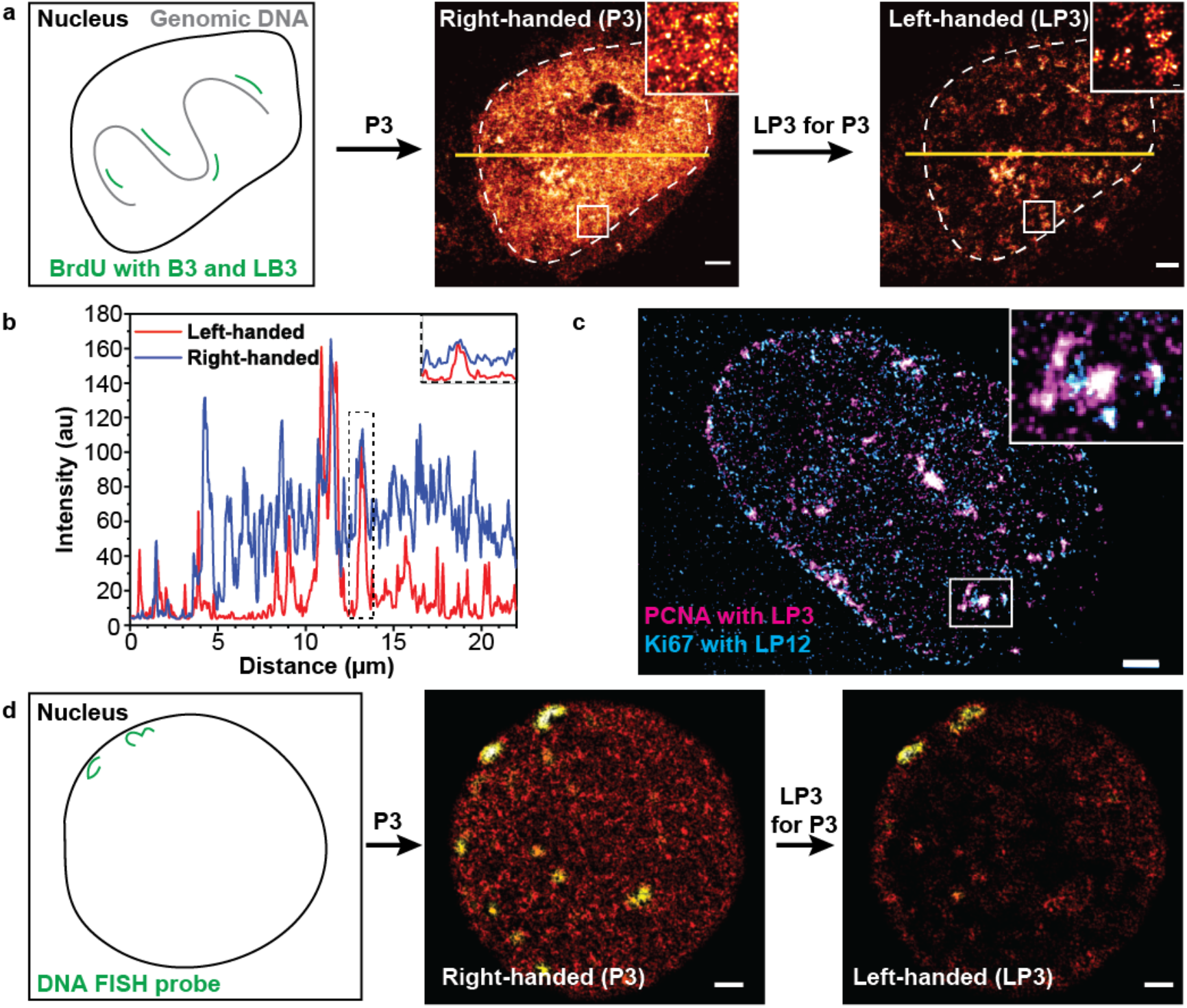
Comparison of L-DNA-PAINT and R-DNA-PAINT for nuclear targets. **a)** Experimental setup for the imaging of DNA replication foci via BrdU incorporation into chromosomes. Cells are stained with a 1:1 mix of B3- and LB3-coupled antibodies against BrdU and undergo both R-DNA-PAINT and L-DNA-PAINT imaging. **b)** Intensity profiles of a cross-section of the BrdU stained cell in **a)**, depicted in yellow. The inset shows that P3 and LP3 stained cells show the same local maxima, but the background intensity in the LP3 stained cell is significantly reduced. **c)** L-DNA-PAINT multiplexing experiment in a Hela cell nucleus stained with a GFP nanobody coupled to LB3 directed against overexpressed GFP-PCNA and Ki67 labeled with an antibody linked to LB12. **d)** L-DNA-PAINT FISH experiment on HEK293T cells harboring two copies of an integrated HHV-6A genome. The integrated viral DNA was labeled with a HHV-6A specific probe, coupled to B3 and LB3 in an equimolar ratio. Subsequent R-DNA-PAINT and L-DNA-PAINT experiments facilitated the detection of the two viral DNA loci, as detected by FISH (Supplementary Figure 2). R-DNA-PAINT via P3 resulted in an increased amount of background localizations hampering the detection of specific FISH loci. Scale bars are 2 μm.

An important advantage of DNA-PAINT is the ability to visualize multiple targets in a sample through multiplexing, avoiding chromatic aberration. When we performed L-DNA PAINT against immune-stained PCNA and Ki67 in the nuclei of HeLa cells using different L-DNA imager-binder pairs (LP3-LB3 and LP12-LB12, respectively), we found discrete staining for both molecules consistent with their respective reported localization in the nucleus (Figure 3c).

To further validate our results, we performed fluorescence *in situ* hybridization (FISH), the most frequently-used assay to investigate genomic localization, coupled with our DNA-PAINT approach. FISH was performed on cells infected with Human Herpesvirus 6A (HHV-6A), a ubiquitous beta-herpesvirus that integrates its genome into the telomeres of the host chromosomes (Supp. Fig. 2)^15^. We used previously characterized HEK293T cells harboring two copies of the integrated HHV-6 to test the specificity of L-DNA PAINT technique in the nucleus. Viral DNA was labeled with an HHV-6A specific probe as described previously^16^, which was then coupled in equimolar ration with an L-DNA and an R-DNA binder (LB3 and B3). When we then added P3 imager strands for R-DNA-PAINT, we found strong nuclear background as shown above. On the other hand, subsequent imaging with LP3 after washout resulted in discreet, super-resolved staining of the HHV-6 integration sites (Figure 3d).

Taken together, we demonstrated that L-DNA PAINT has the same specificity, resolution and multiplexing capabilities as traditional R-DNA PAINT. Strikingly, L-DNA PAINT has a drastically lower false detection rate and allows the investigation of structures and events of cellular DNA in the nucleus. This background likely stems from unspecific hybridization events of R-DNA oligomer imagers with endogenous DNA or RNA, as RNA can dimerize with DNA oligomers as well^17^ and is very abundant in cells and nuclei. Superresolution microscopy of the functional organization of chromosomal DNA and nuclear domains are an important frontier in cell biology^18–20^ and, L-DNA PAINT will enable superior investigations of the 3D-genome at length scales relevant for the molecular machineries involved in gene activity, DNA-Repair, splicing and folding, to name a few^12^ in resting and dividing cells, both in eukaryotic and prokaryotic organisms. As such, we suggest L-DNA PAINT as the SMLM method of choice to visualize and quantify DNA-associated molecules with nanoscale resolution specifically and nuclear structures in general.

## Acknowledgements

This study was supported by the ERC starting grant Stg 677673 awarded to BBK and Deutsche Forschungsgemeinschaft (DFG, German Research Foundation) - Project Number 278001972 - TRR 186 to HE.

## Author Contributions

HJG and HE developed the initial workflow, conceived the project and established the L-DNA PAINT approach. GA conducted the FISH experiments. VF synthesized and labeled the left-handed nanobody and secondary antibody. HJG acquired and analyzed the data. HJG and HE wrote the manuscript. GA and BBK edited the manuscript. BBK offered guidance and resources for the project. HE supervised the project. All authors have reviewed and approved the manuscript.

## Methods

Methods can be found online

## Supplementary Data

**Supplementary Figure 1:**
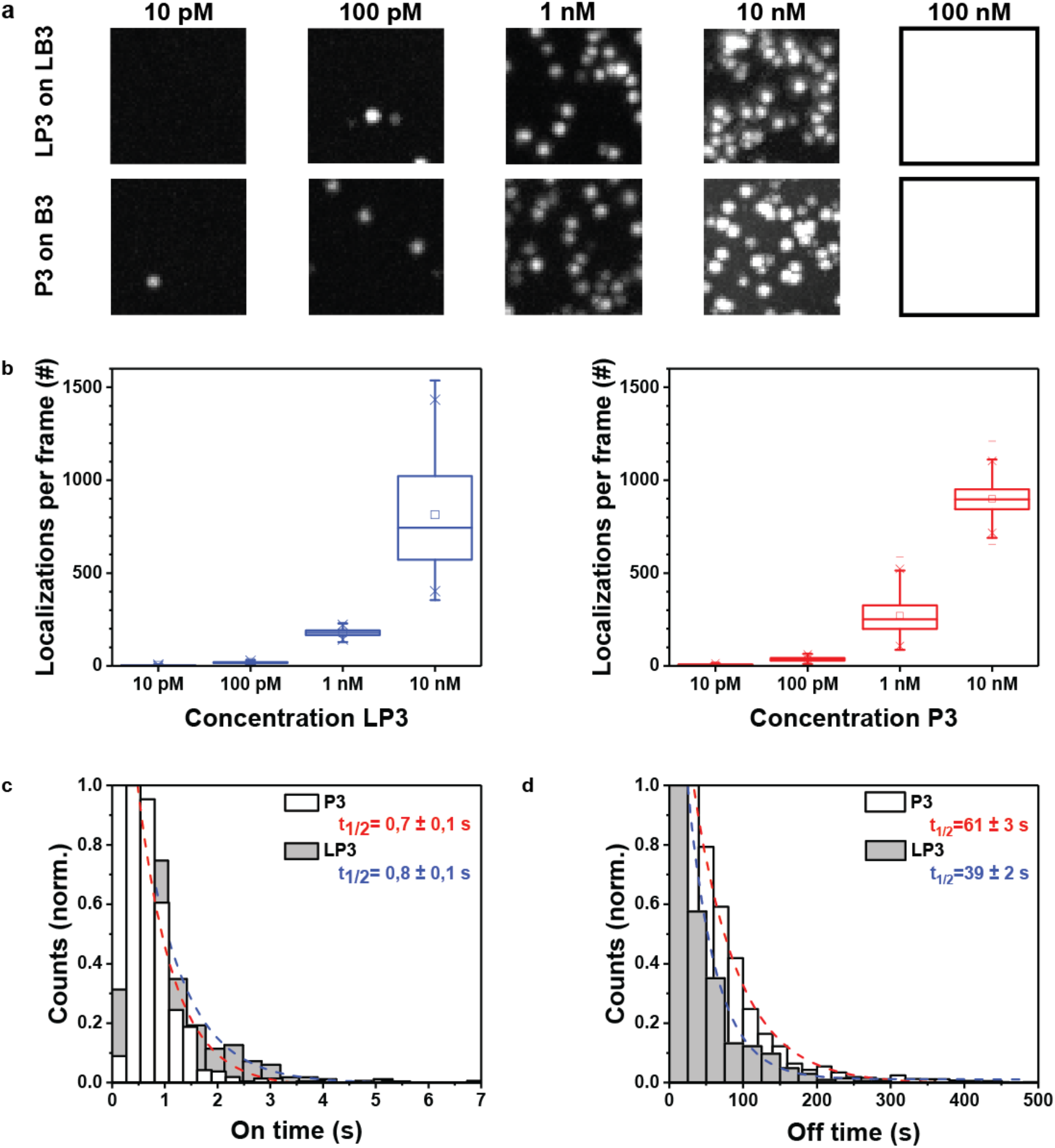
Kinetics of L-DNA and R-DNA PAINT. **a)** Titration experiment of L-DNA imagers (LP3) on L-DNA binders (LB3) and R-DNA imagers (P3) on R-DNA binders (LB3). Single molecule images are 5 μm × 5 μm. **b)** Plot of total number of localizations per frame in the titration experiment was quantified for L- and R-DNA and found to be very similar. No single molecules were detected for 10 nM of imagers and as such excluded from these graphs. **c)** Fluorescence on times as a result of imager-binder hybridization were extracted from the titration experiments. The histogram represents the distribution of on times for L-DNA and R-DNA and was fitted with a single exponential, yielding a half time of 0.8 ± 0.1 s and 0.7 ± 0.1 s, respectively. **d)** The time between subsequent imager-binder hybridization event is defined as the off time. For 1 nM of imager, the off times were found to be 39 ± 2 s and 61 ± 3 s for L- and R-DNA, respectively.

**Supplementary Figure 2:**
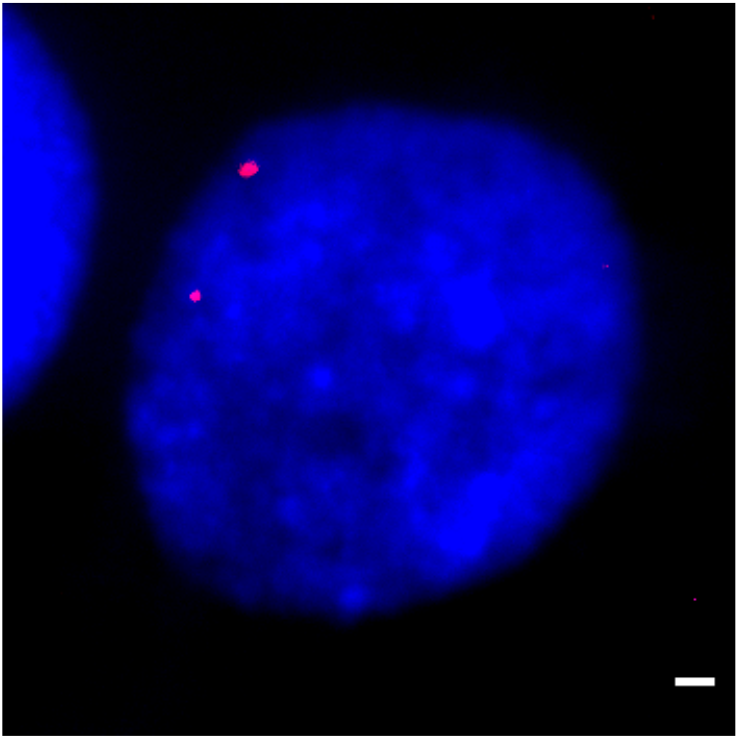
Detection of integrated HHV-6A genomes by FISH. FISH was performed on HEK293T cells harboring two copies of an integrated HHV-6A genome as described previously^16^. The two viral integration sites were detected in the cell nuclei using HHV-6A specific probes by FISH, matching the detection using the DNA-PAINT approach. Scale bar is 2μm.

## Online Methods

### Reagents

All DNA oligomer nucleotides were synthesized from biomers.net. Binder sequences contained a 5’ biotin or 5’ azid modification and imager sequences were conjugated to an Atto655 fluorophore on the 3’ end. In this paper, binder sequences (L)B3 (TTTCTTCATTA) and LB12 (TTAGTTAGAGC) and imager sequences (L)P3 (GTAATGAAGA) and LP12 (GCTCTAACT) were used. Binder oligomers were conjugated to the secondary donkey anti-mouse antibody (Jackson ImmunoResearch Europe Ltd, Cat. Nr. 715-005-150) and GFP nanobody as described previously^1^.

### *In vitro* L-DNA and R-DNA binding assay

Labtek chambers (Ibidi) were cleaned by sonication in 1M KOH. Then, 0.8 mg/mL bovine serum albumin (BSA) and 0.2 mg/mL biotinylated-BSA was added in 10 mM Tris-HCl buffer (pH = 7.5) supplemented with 100 mM NaCl and 0.05% Tween-20. The sample was incubated with 0.2 mg/mL Streptavidin (Sigma Aldrich, Cat. Nr. 85878) to allow binding to the biotinylated BSA on the surface. Afterwards, 500 pM biotinylated LB3 and B3 oligomers were immobilized on the streptavidin in a 5 mM Tris-HCl buffer (pH = 8) supplemented with 10 mM MgCl_2_, 1 mM EDTA and 0.05% Tween-20. Then, the sample was imaged by titrating (ranging from 10 pM to 100 nM) in LP3 or P3 in PBS buffer (pH = 8) supplemented with 500 mM NaCl.

### DNA-PAINT on microtubules in Hela cells

Hela cells were plated on 18mm round coverslips and grown in Dulbecco’s modified Eagle’s medium (DMEM, Life Tech) supplemented with 10% fetal calf serum and 1% GlutaMax (ThermoFisher) at 37C in a 5% CO_2_ humidified incubator. Prior to fixation, the cells were washed 3x with PEM buffer (0.1 M Pipes (pH = 6.95), 2 mM EGTA, and 1 mM MgSO_4_) at 37 C, and permeabilized for 30 s in BRB80 buffer (80 mM Pipes (pH = 6.8), 1 mM MgCl_2_, and 10 mM EGTA) supplemented with 0.5 % Triton X-100. Afterwards, the cells were fixed in 3.2 % paraformaldehyde and 0.1 % glutaraldehyde in BRB80 buffer. Subsequently, the fixation was quenched with 1 mg/mL NaBH_4_ in PBS. Then, the sample was blocked and further permeabilized with 4 % goat serum, 1 % BSA and 0.1 % Triton X-100 in PEM buffer and subsequently with ImageIT (Thermo Fisher Scientific, Inc, Cat Nr I36933). Afterwards, the sample was stained with monoclonal mouse anti-alpha-tubulin antibody (1:1000; Sigma T5618) and subsequently with goat biotinylated anti-mouse antibody (1:200; Polyclonal RU0, BD Pharmingen). Then, 0.2 mg/mL streptavidin (Sigma Aldrich, Cat. Nr. 85878) was added in PBS. Subsequently, 500 pM of the same biotinylated LB3 or B3, as used for the *in vitro* binding assay, were added in PEM buffer. Then the sample was imaged with 500 pM L- or R- imager DNA oligomer’s with Atto647N in PBS supplemented with 500 mM NaCl (also used for the *in vitro* binding assay).

### R- and L-DNA oligomer binding in the nucleus of Hela cells

Hela cells were plated as described above. Then, the sample was washed 1x with PBS and fixed in 4% PFA in PBS. Subsequently, the sample was washed and quenched with 50 mM NH_4_Cl in PBS. Afterwards, the sample was blocked and further permeabilized with 4 % goat serum, 1 % BSA and 0.1 % Triton X-100 in PBS buffer and subsequently incubated with ImageIT (Thermo Fisher Scientific, Inc, Cat Nr I36933). Then the sample was imaged with 500 pM LP3 and P3 in PBS supplemented with 500 mM NaCl (also used for the *in vitro* binding assay).

### DNA-PAINT on BrdU incorporation sites in Hela cells

Hela cells were plated as described above. To stain replication sites, the growth medium was supplemented with 10 μM BrdU for 1 hour and then exchanged for normal growth medium for 1 hour. The cells were fixed in 2% PFA in PBS. Then, the cells were permeabilized in 0.2 % Triton X-100 in PBS. Afterwards, the genomic DNA was denatured by 2 M HCl in PBS. Subsequently, the sample was blocked with 1x BrdU blocking solution (Invitrogen, Cat Nr. 00-4952-52) and stained with anti-BrdU antibody (1 μg/mL; clone BU20A, Invitrogen, Cat Nr. 13-5071-63) in 1x BrdU blocking solution overnight. The sample was stained with streptavidin (0.2 mg/mL; Sigma Aldrich, Cat. Nr. 85878) in PBS and then 1 nM of biotinylated LB3 and B3 (as used for the *in vitro* binding assay) were added. The sample was imaged with 1 nM L- or R-DNA PAINT imagers.

### L-DNA-PAINT on PCNA and Ki67 in the same Hela cell

Hela cells were transfected with GFP-PCNA (Addgene, Plasmid #105977^2^) by an electroporation method (Neon Transfection System, Thermo Fisher Scientific) and plated on 18 mm round coverslips and grown in growth medium. The sample was fixed in 4% PFA in PBS and quenched with 50 mM NH_4_Cl in PBS. Afterwards, the sample was blocked and permeabilized with 4 % goat serum, 1 % BSA and 0.1 % Triton X-100 in PBS and subsequently with ImageIT (Thermo Fisher Scientific, Inc, Cat Nr I36933). The sample was stained with Ki67 (1:100; Clone B56, BD Pharmigen, Cat. Nr. 556003) and afterwards with GFP nanobody coupled to LB3 (1:100; generated and labeled in house) and secondary donkey anti-mouse coupled to LB12 (1:100). The sample was imaged in 100 pM LP3 and 1 nM LP12.

### DNA-PAINT on HEK293T cells stained with Fluorescence *in situ* hybridization (FISH)

The integrated HHV-6A genome was detected by FISH as described previously^3^ with the following modifications^4–6^. Biotin labelled HHV-6A probes were generated using the HHV-6A BAC (strain U1102) and the Biotin-High Prime kit (Sigma-Aldrich, St. Louis, MO, USA). For the classical FISH technique, probes were detected using Cy3-Streptavidin (1:1000; GE healthcare, Chicaco, IL, USA) and DNA was counter stained with DAPI for 10 min (1:3000; Biolegend, San Diego, CA, USA). For the FISH technique coupled with the DNA-PAINT approach, probes were stained using streptavidin (0.2 mg/mL; Sigma-Aldrich, Cat. Nr. 85878) and an equimolar (2 nM) of B3 and LB3, and detected using 2 nM P3 and LP3.

### Microscopy

Images were acquired with a Vutara 352 super-resolution microscope (Bruker) equipped with a Hamamatsu ORCA Flash4.0 sCMOS for super-resolution imaging and a 60× oil immersion TIRF objective with numerical aperture 1.49 (Olympus). Data was required with TIRF/HILO-illumination at a laser power density of ~2.5 kW/cm^2^ using a 639 nm laser. Images were typically collected at 300 ms acquisition time and typically 5000 images were collected for the *in vitro* binding assay and 15000 images were used to reconstruct the super-resolution composites. Data was analyzed using the Picasso software (Fig. 1-3c) or by a representation of the pair correlation of all localizations by Vutara’s SRX software.

